# Rapid Construction of a Whole-genome Transposon Insertion Collection for *Shewanella oneidensis* by Knockout Sudoku

**DOI:** 10.1101/044768

**Authors:** Michael Baym, Lev Shaket, Isao A. Anzai, Oluwakemi Adesina, Buz Barstow

## Abstract

Whole-genome knockout collections are invaluable for connecting gene sequence to function, yet traditionally they have needed an extraordinary technical effort to construct. Knockout Sudoku is a new method for directing the construction and purification of a curated whole-genome collection of singlegene disruption mutants generated by transposon mutagenesis. Using a simple combinatorial pooling scheme, a highly oversampled collection of transposon mutants can be condensed into a next-generation sequencing library in a single day. The identities of the mutants in the collection are then solved by a predictive algorithm based on Bayesian inference, allowing for rapid curation and validation. Starting from a progenitor collection of 39,918 transposon mutants, we compiled a quality-controlled knockout collection of the electroactive microbe *Shewanella oneidensis* MR–1 containing representatives for 3,667 genes. High-throughput kinetic measurements on this collection provide a comprehensive view of multiple extracellular electron transfer pathways operating in parallel.

## Introduction

*Shewanella oneidensis* MR-1 is one of the archetypal members of the electroactive microbes^1^. Members of this class are capable of transferring electrons to, from or between metabolism and a remarkable variety of external substrates, ranging from metal deposits to electrodes^1^. The *Shewanellae* serve as model organisms for understanding the role of microbes in the biogeochemical cycling of metals^2^ and carbon^3^ as well as the geochemical evolution of the Earth^4,5^. *S. oneidensis* possesses one of the largest complements of cytochrome electron transfer proteins^6^, including the *mtr* extracellular electron transfer (EET) operon that encodes a set of proteins responsible for most of the electron flux from metabolism to external electron acceptors^3^. Its extensive complement of cytochromes provides *S. oneidensis* with extraordinary respiratory flexibility^7^ and give it potential applications in environmental remediation^2,4^, nuclear stewardship^8^ and sustainable energy^9,10^, while its role in the cryptic cycling of sulfur may have consequences for the fate of CO_2_ sequestered in deep aquifers^11^. However, despite intense research, the functions of over half of the genes in the *S. oneidensis* genome remain unknown^12^ and fundamental discoveries about the nature of the organism continue to be made^11,13^. Although the centrality of the MtrA, MtrB and MtrC cytochromes in EET is well established^14^, debate about the full complement of proteins used in EET, particularly in the transfer of electrons from the inner membrane across the periplasmic gap to the outer membrane continues^15-17^. There remains an ongoing need for the continued development of new tools for the genetic manipulation and characterization of *S. oneidensis*, as well as other electroactive microbes and esoteric microorganisms^1,7,18^.

Aside from the whole genome sequence, a single-gene knockout collection is arguably one of the most valuable genetic tools for any organism. Transposon mutagenesis allows for the straightforward creation of extremely large single-gene disruption mutant libraries for a wide variety of microorganisms^19,20^. Recently developed techniques that use massively parallel sequencing of pooled transposon insertion mutant libraries^21-23^ have made dramatic advances in the characterization of genes that can be selected for fitness contributions^24^ to species including in *S. oneidensis*^*12,25*^. Nevertheless, clonally isolated collections of mutants remain a critically important tool for the characterization of phenotypes such as virulence factors^26,27^, secondary and cryptic metabolite production^28^ and specific behaviors synonymous with *Shewanellae*, such as biofilm formation^29,30^ and EET^31-33^. However, the random nature of transposon mutagenesis requires that non-annotated collections be extremely large to ensure coverage of each nonessential gene in the genome^34^ and does not facilitate testing of specific predictions of genotype-phenotype associations^35^. The utility of comprehensive, non-redundant collections is hard to overstate: in the 15 years since the release of the first version of the *S. cerevisiae* gene deletion collection, it was used in over 1,000 genome wide experimental screens^36^. While curated, annotated collections remain the gold standard, their construction either by Sanger sequencing of extremely large libraries made by random transposon insertion and subsequent condensation^27,37,38^ or by targeted deletion^34,39-42^ is an extraordinary technical endeavor and as a result, only a handful of these types exist^21^.

The emergence of massively parallel sequencing methods has led to an explosion of genomic data in the past decade. In addition, recent advances in robotic combinatorial pooling methods^24,43^ have allowed the annotation of arrayed collections of clonally isolated samples through a single massively parallel sequencing experiment. This exciting development considerably reduces the sequencing costs of annotation and has facilitated the construction of a growing number of curated collections of transposon insertion mutants of pathogens and pathogen surrogates^35,44,45^. Although such methods allow for robust sequence analysis and mapping, the hardware needed is not yet widely deployable. This has inspired developments in simple combinatorial pooling methods that do not rely on robotics^46^. However, the lack of complexity in the combinatorial pools must be compensated by sophisticated algorithms for the disambiguation of sequencing data.

Here, we describe an easily implemented, rapid and generalizable method: Knockout Sudoku, that is capable of automatically cataloging extremely large transposon insertion mutant collections through a simple four-dimensional combinatorial pooling scheme and analysis of data from a single massively parallel sequencing experiment (**Fig. 1; Supplementary Fig. 1**). A Bayesian inference algorithm that uses self-consistency within the dataset is used to disambiguate the location of mutants appearing at multiple locations in the collection. Using this method, we catalogued a 39,918 member transposon insertion collection for *S. oneidensis* that is suitably large to ensure disruption of a large fraction of the nonessential open reading frames in the genome. This catalog was used to predict the contents of each well in this progenitor collection and algorithmically guide the re-array and colony purification of a nonredundant set of mutants. This collection was then independently validated through a second round of combinatorial pooling, sequencing, and orthogonal sequence analysis. In total, the quality-controlled collection comprises isolated mutants for 3,282 genes, which was later expanded to include an additional 385 genes. Furthermore, we functionally validated this collection through a well established colorimetric assay for EET that has been applied to traditional transposon collections of *S. oneidensis*. Using high-throughput photography, time courses of EET were captured nearly simultaneously for every mutant in the collection, recapitulating previous discoveries in a single experiment as well as verifying the functions of associated genes. With the wealth of sequencing data now available for the *Shewanellae*^47^ and other esoteric organisms, the Knockout Sudoku method provides an exciting avenue for comparative genomics and systems biology.

**Figure 1:**
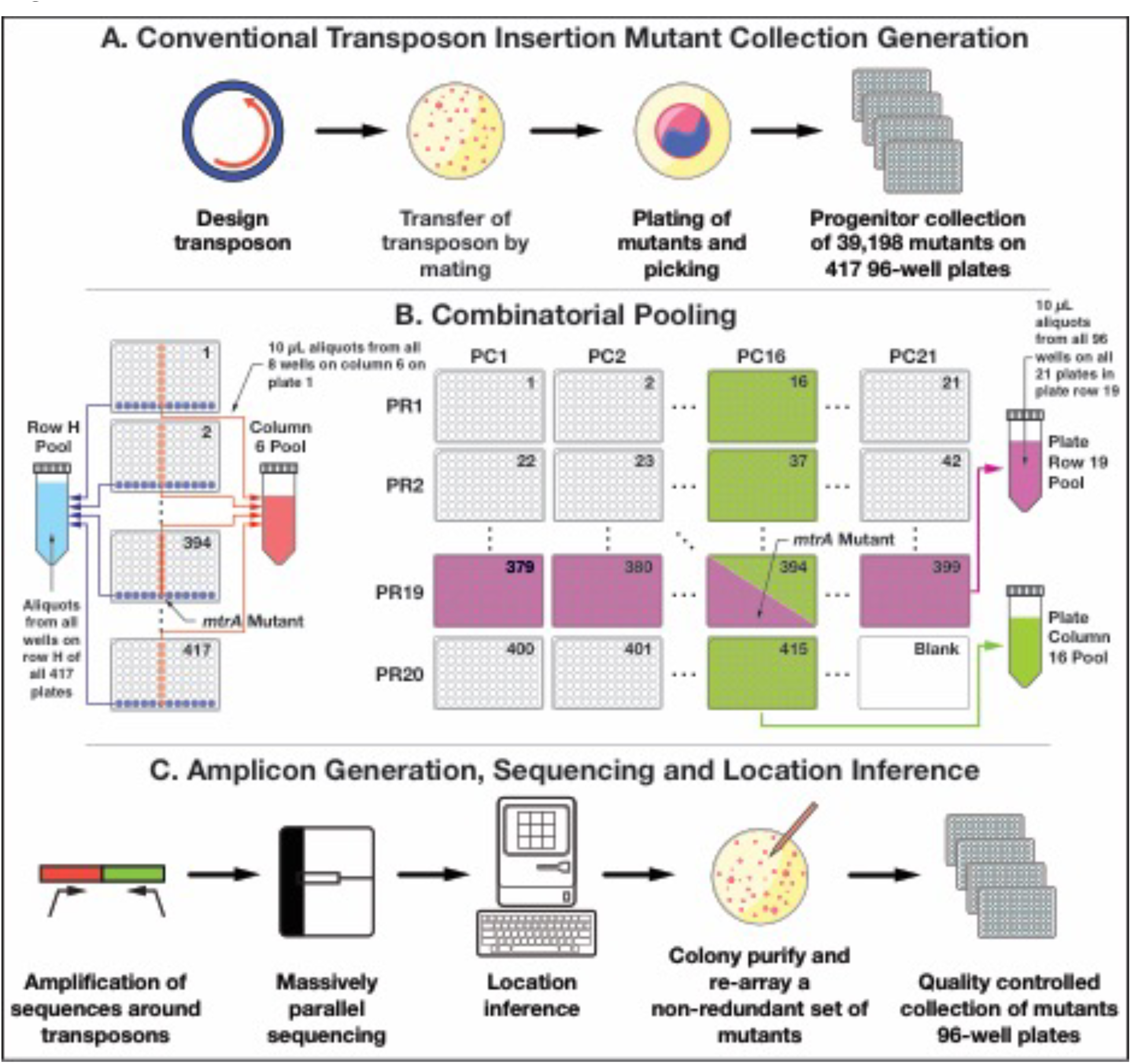
Workflow for creation of the quality-controlled *S. oneidensis* Sudoku collection by Knockout Sudoku.

## Results

### Construction of the Progenitor Collection

The first steps of Knockout Sudoku follow those of conventional transposon-based collection generation (**Fig. 1A**). We constructed a transposon insertion plasmid based on pMiniHimar^19^ containing flp-recombinase sites to permit later excision of the antibiotic resistance cassette. The transposon plasmid was delivered to *S. oneidensis* by conjugation with *E. coli* WM3064^48^. The appropriate size of the progenitor collection was estimated by two methods. An analytical Poisson-based model, which is only a function of the number of non-essential genes, provides a baseline minimum of ≈ 20,000 mutants for complete coverage of the genome (**Fig. 2, dashed lines**), while a Monte Carlo numerical simulation of transposon insertion that takes into account gene location and essentiality data estimates that we would need at least twice this many (**Fig. 2, solid lines; Online Methods 1**). Based upon the more conservative of these estimates, we picked a progenitor collection of 39,198 *S. oneidensis* transposon insertion mutants over the course of two days and arrayed them into 417 96-well plates (**Online Methods 2**).

**Figure 2:**
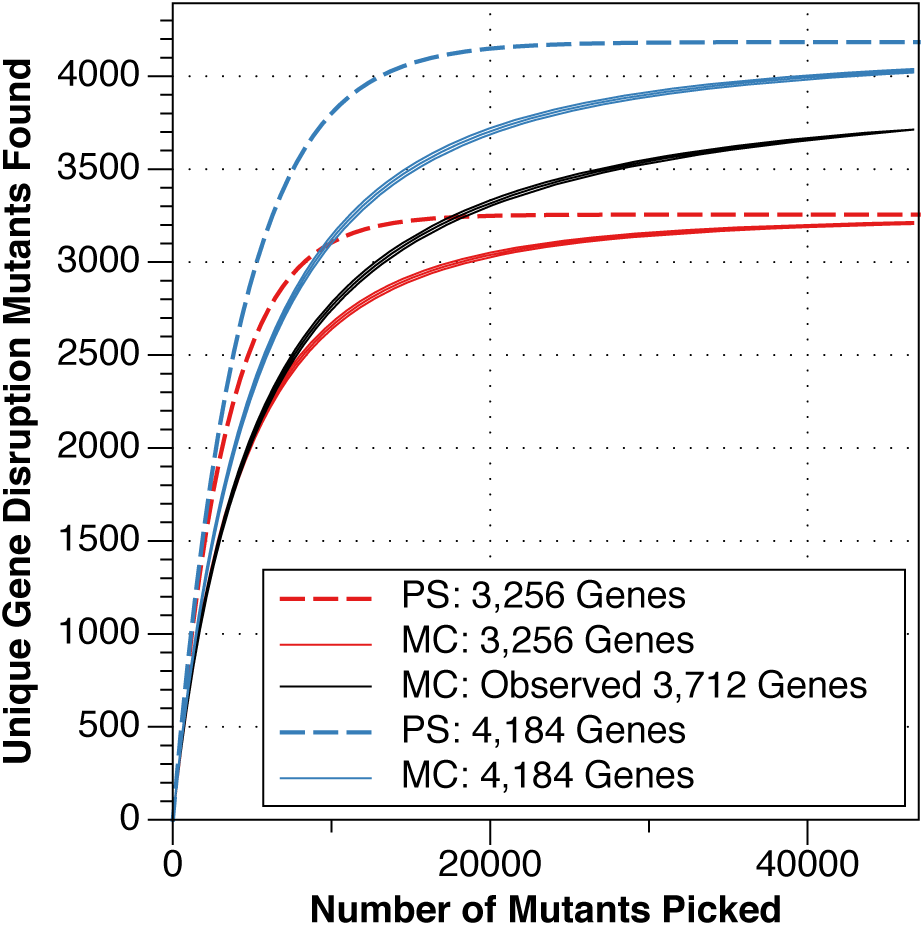
Comparison of analytical (PS, dashed lines) and numerical (MC, solid lines) estimates of the number of non-essential genes disrupted relative to transposon insertion mutant collection size for an upper (4,184 out of 4,587 loci, blue curves) and lower (3,256 genes, red curves) estimate of the *S. oneidensis* non-essential gene set, alongside a numerical estimate of disrupted gene count using a random drawing without replacement from the observed set of transposon insertion mutants (black curves). The center of the solid curves is the mean value of the unique gene disruption count from 1,000 simulations while the upper and lower curves around represent 2 standard deviations around this mean.

### Combinatorial Pooling Process for Knockout Sudoku

The goal of combinatorial pooling is to combine a collection of many samples into a single massively parallel sequencing experiment and then reconstruct the physical locations of each unique sequence. This is made possible by distributing aliquots of each sample to a set of pools determined by its location in the collection, extracting the essential sequence information from that pool, and adding a DNA barcode unique to the pool that is later used to map sequences to physical locations in the collection.

For the Knockout Sudoku method, we developed a 4-dimensional combinatorial pooling scheme that can be easily performed with multi-channel pipettors while minimizing sample preparation costs (**Fig. 1B**). All 417 plates in the collection were assigned to a position within a virtual 20 × 21 grid, giving each plate two coordinates: a plate-row (PR) and a plate-column (PC) (**Fig. 1B**). The assignment of 2 coordinates (rather than 1) per plate allows the cost of sequencing library construction to grow only with the square root of the number of plates. Aliquots of culture from each well in the collection were dispatched to 4 pools that uniquely corresponded to the address of the well within the plate grid and the individual plate. For example, a mutant with a disruption in the *mtrA* cytochrome that was located in well H6 on plate 394 was dispatched to the Row H, Column 6, PR 19, and PC 16 pools (**Fig. 1B**). In total, we filled 61 address pools (20 plate-row × 21 plate-column pools, 8 row × 12 column pools). The entire progenitor collection was pooled and cryopreserved in a single day using a 96-channel pipettor and a team of 5 people (**Online Methods 3**).

The address pools were used to generate 61 barcoded amplicon libraries that encoded the genomic locations of the transposons present within each pool. A nested PCR reaction^49^ was used to amplify the transposon insertion site and add custom sequencer-compatible sequencing-primer-binding sites and flowcell binding sequences (**Supplementary Fig. 2; Online Methods 4**). The standard Illumina index sequence was substituted for a custom barcode sequence that is unique to each pool. The libraries were then combined and single-end sequenced from the transposon side to at least 40 bases past the junction on 2 lanes of an Illumina HiSeq 2500 (**Online Methods 4**).

### Location of Transposon Insertion Mutants in the Progenitor Collection

To locate mutants within the progenitor collection, the sequencing dataset was parsed to a set of putative transposon insertion locations and pool address coordinates. **Fig. 3A** shows an annotated read from an amplicon produced by the *mtrA* disruption mutant highlighted in **Fig. 1**. A flow chart of the sequence analysis workflow can be found in **Supplementary Fig. 1**. Overall, of the 146,128,068 initial reads generated by Illumina, 120,085,483 aligned to genomic locations and contained valid barcode and transposon sequences (**Online Methods 5; Supplementary Fig. 1, algorithm 5**). For each unique genomic location, we counted the number of reads with a given barcode to construct a pool presence table that was used to deduce the location of mutants within the collection. Of the 83,380 unique entries in the pool presence table, we find that roughly half correspond to low-frequency amplification events that are unlikely to be due to actual transposon insertions in the genome: 19,723 entries are associated with only a single read, while another 16,754 were associated with 2-5 total reads. Such low-frequency events were systematically filtered by requiring a threshold number of reads per coordinate (**Supplementary Fig. 3**).

**Figure 3:**
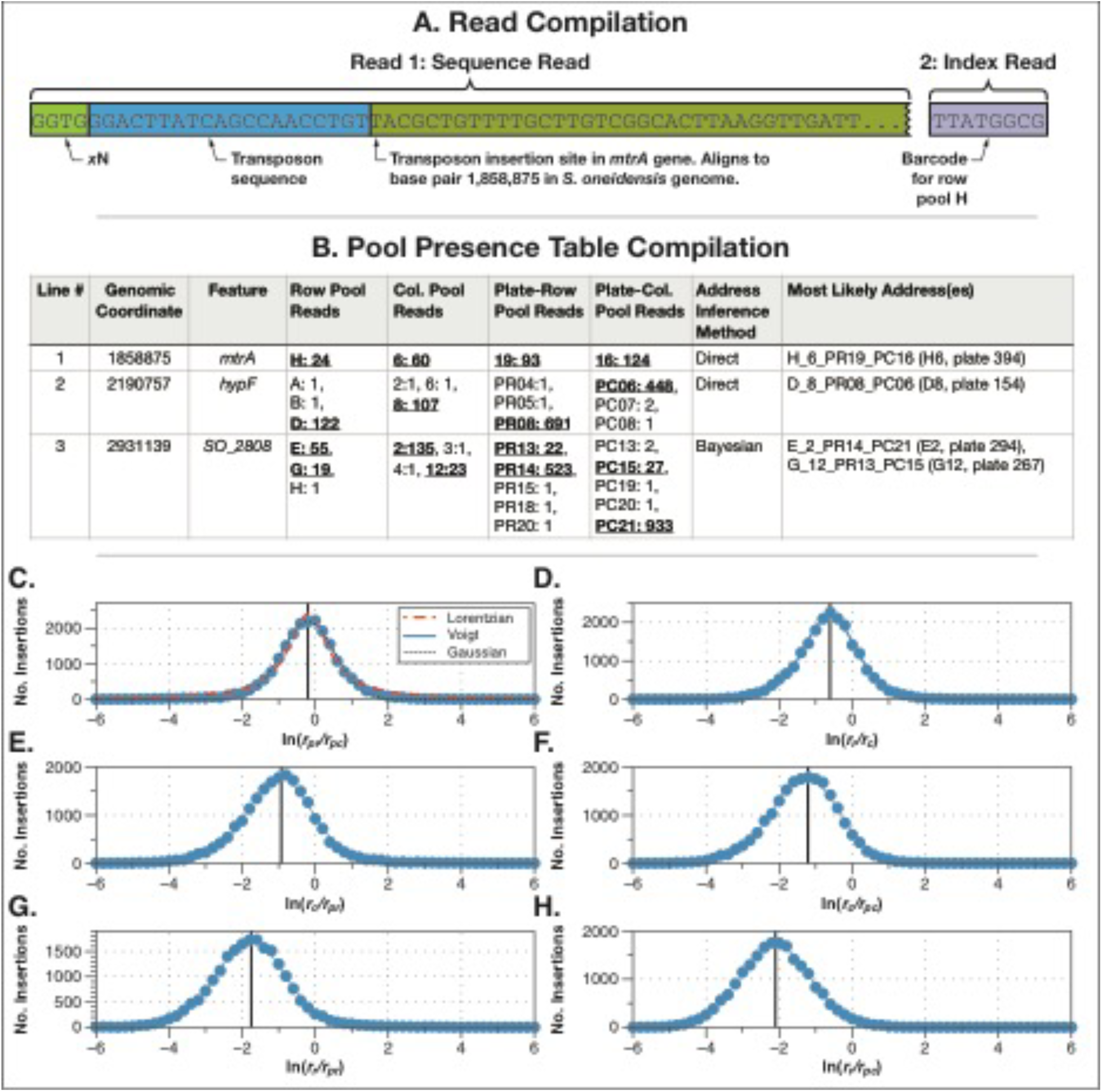
Knockout Sudoku sequence data analysis. **A:** Reduction of read data to transposon insertion coordinates and pool barcode identities. **B:** Compilation of a pool presence table that summarizes the number of reads aligning to a given genomic coordinate with a given pool barcode. Validation of predictions from lines 2 and 3 by Sanger sequencing can be found in **Supplementary Table 2. C-H:** Distribution of read count ratios for pool presence table lines that unambiguously map to single addresses in the progenitor collection ( **line 1**). Distributions of **C:** plate-row (pr) to plate-column (pc); **D:** row (r) to column (c); **E:** column to plate-row; **F:** column to plate-column; **G:** row to plate-row; **H:** row to plate-column read count ratios. The center of the distribution is marked with a vertical line, which moves to lower values as the relative amount of template per mutant is reduced in the amplicon generation reaction (**Supplementary Fig. 2; Online Methods 4**). **C** shows a comparison of log-Voigt, -Gaussian and - Lorentzian fits.

A threshold of 5 read counts per coordinate leads to a stable solution of the pool presence table with a physically reasonable number of entries (39,588) that have one or more coordinates that can be used for mapping. Three sample lines are shown in **Fig. 3B**. Here, line 1 corresponds to the *mtrA* disruption mutant highlighted in **Fig. 1B** and simply maps to well H6 on plate 394.

In the final pool presence table, we find that the majority of the entries map to addresses in the progenitor collection in a simple manner: 20,973 map in a one-to-one fashion to single addresses, as in the *mtrA* example (**Fig. 3B, line 1**), while 5,131 entries map unambiguously to multiple addresses. The second type of entry can arise if, for example, the same colony was picked into 2 wells in the same row of a single plate. The remaining entries either had too little or too much information to be immediately mapped: 8,424 did not have a complete set of coordinates, whereas 5,060 had presence in multiple coordinates that could be combined in multiple ways. For example, the coordinates shown in **Fig. 3B (line 3)** most likely originate from only 2 addresses (one set that has a high read count and a second that has a low read count) but can form 16 possible combinations.

Internal self-consistency within the sequencing dataset allows for the automatic deduction of the location of mutants that do not simply map to addresses in the progenitor collection (**Online Methods 5**). Mutants that appear at multiple originating addresses can produce coordinates that combine into both the correct addresses and a far greater number of incorrect addresses. However, the frequency of a given amplification event should not vary significantly from pool to pool, and therefore, the ratio of the reads from multiple pools should be fixed around a mean value. We find this intuition to be true using the 20,973 entries in the pool presence table that map to single addresses in the collection. The six ratios of the read counts for the four coordinates (R, C, PR, PC) were computed for every entry to generate histograms (**Fig. 3C-H, blue circles**). We find that logarithms of these distributions are excellently fit by Voigt functions (**Fig. 3C-H, blue curves**). The centers of these distributions move to lower values because the quantity of template per individual mutant used in the amplicon library generation reaction decreases as the number of species in a pool increases (**Fig. 3C-H**). For example the amount of template of an individual species in a row pool reaction should be 8/12^th^ of that same species in a column pool reaction, as each 96-well plate has 8 rows and 12 columns. Additionally, these peaks are shifted to even lower values due to correlation between individual sets of read counts. The log-Voigt functions can then be normalized to an integrated area of 1.0, converting them to a set of probability density functions. This permits us to rate the probability that a given combination of coordinates contains the mutant through Bayesian inference by simply multiplying together the probabilities of the read count ratios for a given combination of coordinates in a proposed address.

In total, we mapped 31,860 mutants to 21,907 singly-occupied and 9,922 multiply-occupied wells in the progenitor collection. The remaining 7,369 wells are a likely home for the mutants for which we have insufficient address data. Histograms of the density of mappable transposons in the progenitor collection across the *S. oneidensis* chromosome and megaplasmid are shown in **Fig. 4** along with a plot showing the location of individual transposons in the vicinity of the *mtr* EET operons. The transposon coverage across the chromosome (61 transposons per 10,000 bp) and megaplasmid (110 per 10,000 bp) is approximately constant. Overall, we were able to find disruption mutants for 3,474 loci in singly-occupied wells while disruption mutants for 238 could only be found in multiply-occupied wells. A complete catalog of the predicted contents of the progenitor collection can be found in **Supplementary Table 1**. The predicted contents of 174 unique members of the progenitor collection were individually verified by Sanger sequencing. Of these predictions, 163 were accurate to within 1 bp, with none more than 5 bp (**Online Methods 6; Supplementary Table 2**).

**Figure 4:**
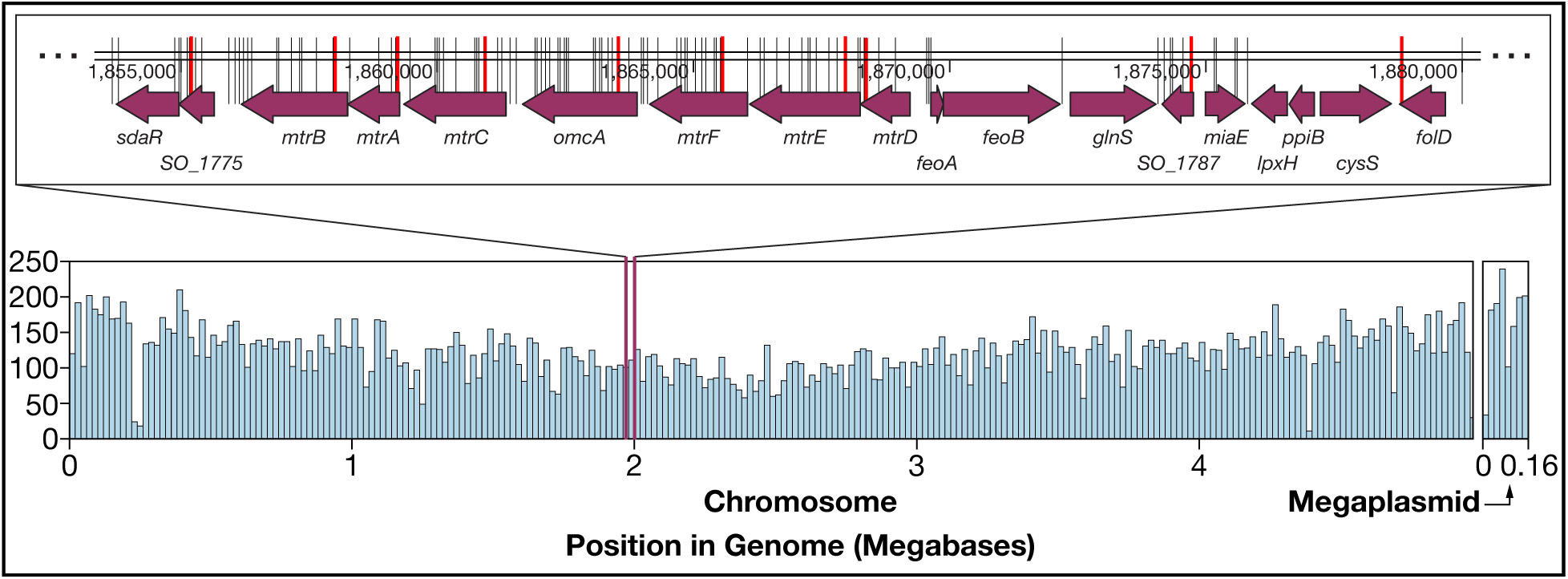
Lower Plot: Histogram showing the density of transposons in the progenitor *S. oneidensis* Sudoku collection (bin width is 20,000 base pairs). **Upper Plot:** Map of locations of transposons in the progenitor collection (black lines) and those chosen for the quality-controlled collection (red lines) in the vicinity of the *mtrABC* and *mtrDEF* extracellular electron transport operons. It is important to note that the chosen mutant was not always the one closest to the translation start, as in a number of cases our selection algorithm (**Online Methods 7**) estimated that the benefits to gene disruption were outweighed by the difficulties of isolation.

### Generation and Sequence Validation of a Quality-Controlled Collection

The progenitor collection catalog was used to assemble a quality-controlled non-redundant collection of 3,630 mutants that disrupt a total of 3,667 genes. Oversampling not only ensures sufficient coverage of the genome but allows us to choose the mutant that best disrupts a given gene. We developed a set of algorithms that identified the closest mutant to the translation start of each gene that was the most easily isolated (**Online Methods 7; Supplementary Fig. 4**). The mutants chosen to represent each of the genes in the *mtr* operons are highlighted in red in **Fig. 4**. We generated a condensed collection by re-array of 2,699 representative mutants from singly-occupied wells, and colony purified representatives for another 999 loci. Based on the predicted content of the well, we picked between 2 and 10 colonies for each of the colony-purified mutants into a series of 96-well plates. In total, this collection comprised 5,653 samples, arrayed on 84 plates. The collection was arranged into a 9 × 10 plate grid, re-pooled and re-sequenced (**Online Methods 8; Supplementary Fig. 4**).

We used an orthogonal sequence validation algorithm to test if the predicted content of each well in the condensed collection was present (**Online Methods 8; Supplementary Fig. 4**). Of the 5,653 mutant identity predictions, 5,202 were correct, a 92% accuracy rate that indicates success in sequence location, initial re-array, colony purification and picking. It is important to note that a finite error rate in re-arraying should be expected: the first generation Keio Collection has a 4% error rate^50^. However, a key advantage of the low cost and convenience of Knockout Sudoku is the ability to immediately diagnose this error. In total, we verified the identity of 3,245 mutants, disrupting 3,282 loci. In addition, we later added 385 mutants, bringing the total of disrupted loci to 3,667. With this extension, the quality-controlled collection represents almost 99% of the disruptable loci identified in the progenitor collection. A complete catalog of the quality-controlled *S. oneidensis* collection can be found in **Supplementary Table 3**.

### Functional Validation of the Quality-Controlled Collection

We screened the quality-controlled *S. oneidensis* mutant collection for EET activity with a well-established colorimetric assay that uses anthraquinone-2,6-disulfonate (AQDS), an analog for humic substances that can act as electron acceptors during anaerobic respiration^31^. AQDS changes color from clear to deep orange when reduced by 2 electrons, allowing visual identification of mutants that are deficient in extracellular reduction capability. A time series of images was captured for every plate in the collection over 40-60 hours under an anaerobic atmosphere and was automatically reduced to a time course for each mutant in the collection (**Figs. 5** and **6; Supplementary Figs. 5** and **6**; **Online Methods 10** and **11**). The linear portion of each time course was fitted to calculate a reduction rate (**Fig. 6D; Supplementary Table 5**).

**Figure 5:**
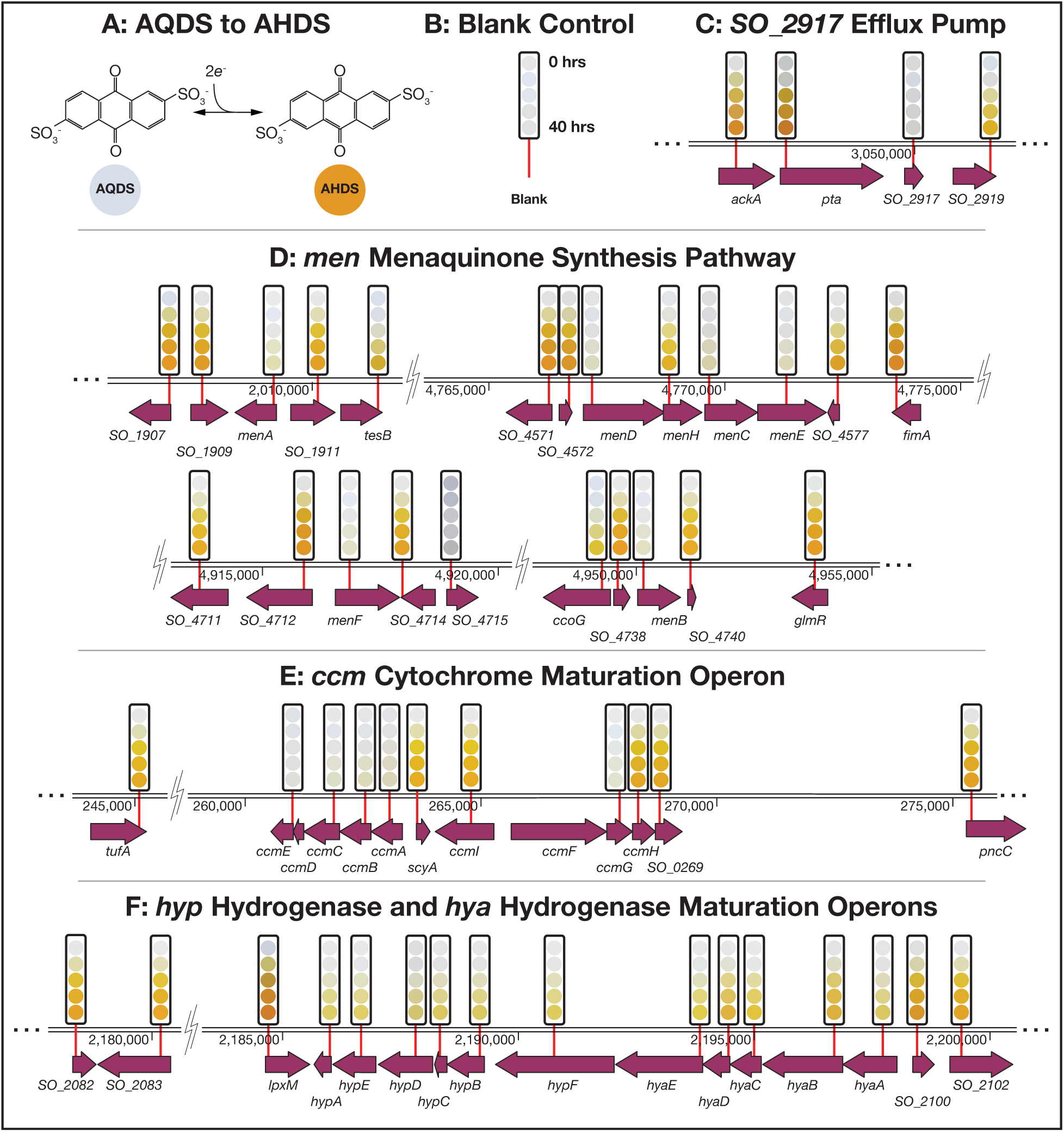
AQDS reduction screen results showing gene disruptions that produce large changes in extracellular electron transfer (EET) ability of *S. oneidensis*. **A:** AQDS redox reaction. AQDS changes color from clear to orange when reduced. **B:** Blank control. **C-F:** The state of the AQDS dye at 10 hour intervals is indicated by a series of colored circles above the location of the transposon chosen to disrupt each gene (**Online Methods 10** and **11**).

**Figure 6:**
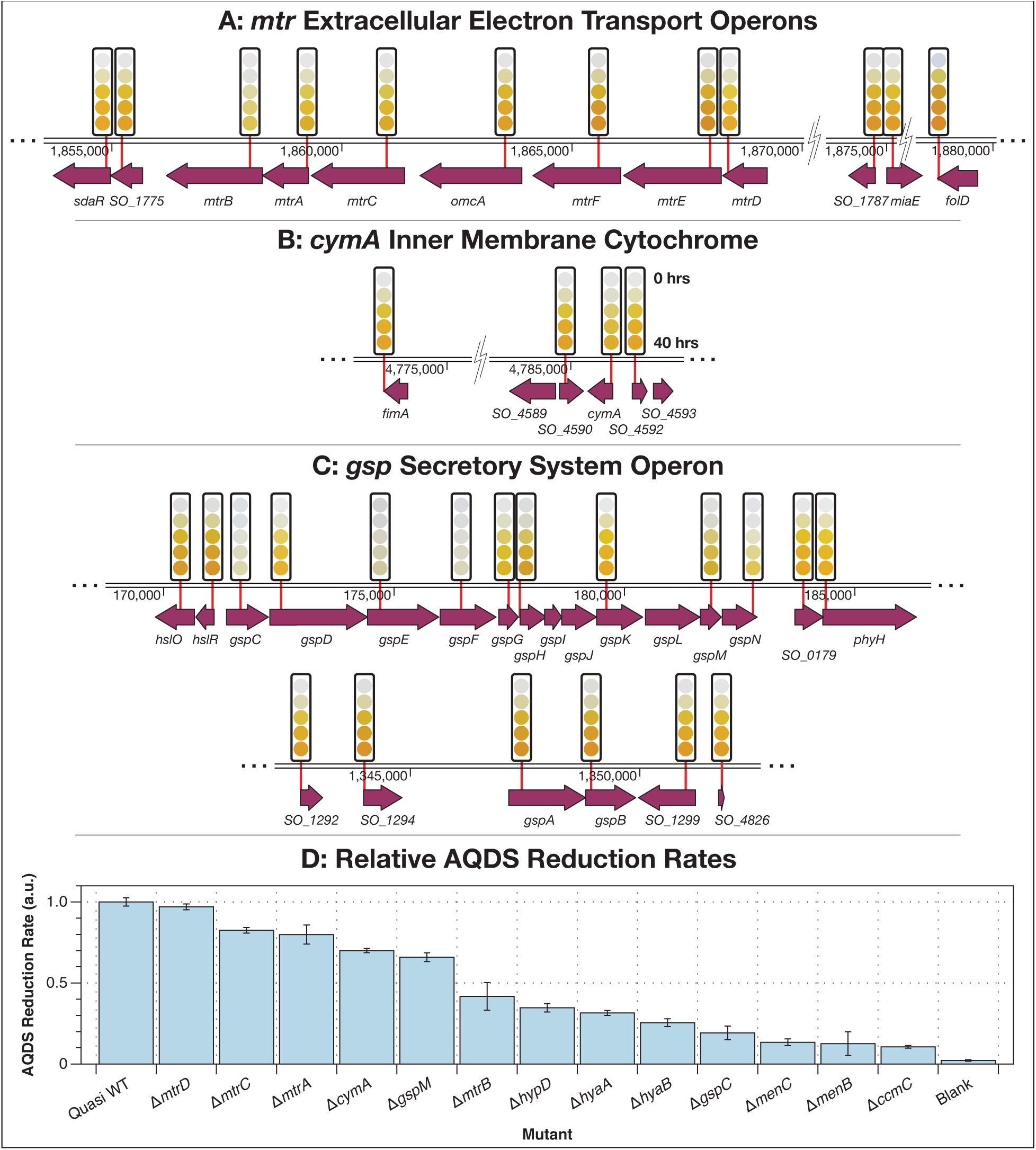
A-C: AQDS reduction screen results showing gene disruptions that produce small but detectable changes in EET activity. **D:** Linear AQDS reduction rates of selected mutants relative to an averaged quasi wild-type (WT) control (**Online Methods 10** and **11**). A full listing of rates for all mutants can be found in **Supplementary Table 5**.

Our screening results include previously identified genes as well as additional associated genes. Mutants with disruptions in the *menC* menaquinone synthesis gene^31^; the tolC-like efflux pump *SO_2917* that moderates AQDS toxicity^51^ and the *ccmC* cytochrome maturation gene^19^ all showed large deficiencies in their ability to reduce AQDS (**Figs. 5C, D and E**). In addition, a dramatic loss of AQDS reduction ability was registered for disruption mutants of the menaquinone synthesis pathway members *menA, B, D, E*, and *F* and the cytochrome maturation operon members *ccmA, B, D, E* and *G* (**Figs. 5D and E**).

Disruption of any of the genes encoding the Hyp NiFe uptake hydrogenase, *hypA, B, C, D* and E, and its maturation factors *hyaA, B, C, D* and *E* leads to significant loss of AQDS reduction rate and completeness over the course of an experiment (**Fig. 5F**). These results lead us to speculate that oxidation of H_2_ is a significant electron source for EET under the conditions of our screen.

In agreement with earlier measurements, disruption of the MtrB outer-membrane porin responsible for supporting the outer-membrane metal reductase MtrC, slows AQDS reduction^51^ to 42% of the quasi wildtype rate but does not completely eliminate it, consistent with reports of the operation of multiple EET pathways^52^. Similarly, disruption of MtrC slows AQDS reduction to ≈ 83% of the quasi wild-type, consistent with earlier reports that this mutation does not completely eliminate soluble and insoluble Fe(III) reduction^53^. Disruption of the paralogous *mtrDEF* operon had no discernible impact on AQDS reduction. Consistent with a role for the Mtr system in AQDS reduction, disruption of the type II secretion system, considerably slows AQDS reduction^54^. However, disruption of CymA, a key distribution point for electrons to a variety of reductases including the Mtr system^14,55^, only slows AQDS reduction to 70%. Most, notably, in contrast to experiments with Fe(III) reduction, disruption of MtrA only slows AQDS reduction to 80% of the quasi wild-type rate^53^.

Disruptions to the *ccm* operon indicate a crucial role for cytochromes, such as those encoded by the *mtr* genes, in electron transfer to AQDS. However, no disruption in any of the genes in the *tor* (TMAO reductase); *sir* (sulfite reductase); and *dms* (DMSO reductase) operons produced any noticeable diminution of AQDS reduction (**Supplementary Fig. 6; Supplementary Table 5**)

## Discussion

The recapitulation and extension of many of the results of multiple earlier studies on EET in *S. oneidensis* demonstrates the basic functionality of the *S. oneidensis* Sudoku collection and gives us confidence that this and future collections made by Knockout Sudoku can be reliably used in a wide range of genetic screens. The conditions of this EET screen, along with high-throughput kinetic measurements, allow discrimination between mutants with different EET rates that may not be apparent to the eye.

The most notable result is the necessity of cytochrome maturation, but the high apparent replaceability of the MtrA, MtrC and CymA cytochromes under these conditions. None of these mutations slow AQDS reduction by more than 35%. In contrast, disruption of MtrB slows AQDS reduction by 60%. These results point to two conclusions: firstly, that multiple cytochrome-dependent EET pathways operate under the conditions of this screen. Secondly, given that disruptions to MtrA, MtrC and CymA produce much smaller rate reductions than the MtrB disruption it is possible that these proteins can be substituted for alternative proteins expressed under the conditions of the screen. Alternatively, the MtrB porin could perform a role in AQDS reduction beyond simply hosting MtrA and C such as allowing AQDS greater access to the cytochromes in the periplasm. This stands in contrast to results on Fe(III) reduction, where elimination of MtrA produces almost complete elimination of EET^53^. The surprisingly small AQDS reduction defect caused by disruption of CymA could suggest the operation of another inner membrane quinone oxidoreductase.

Arrayed, condensed, near-complete knockout collections, like the one generated here, have a number of advantages for genetic screening. Segregation of isolates permits screening for loss-of-function mutations and for phenotypes which do not appreciably change growth rate, neither of which is possible with a pooled collection. The small size of a condensed collection allows much finer-grained experiments, like colorimetric time courses, which would be impractically onerous on a 40,000 member collection. Annotation allows us to examine a subset of a collection and carefully test for slight deviations from normal behavior and confirm the lack of an involvement of any gene in a process.

The full complement of proteins needed to transfer of electrons across the periplasmic gap remains a subject of debate, with some investigators suggesting a role for additional periplasmic proteins. The *S. oneidensis* Sudoku collection, coupled with the high-throughput kinetic screening demonstrated here is a valuable tool for searching for them^16^. Insights gained may provide valuable information for the further performance improvement of EET machinery in engineered heterologous hosts^56,57^.

The wealth of sequencing data on the *Shewanellae*^47^ is illustrative of the growing abundance of such data for groups of closely related species and esoteric microbes. Perhaps now more than ever, new technologies are needed to connect this to the functions of novel microorganisms^23^. Curated whole-genome knockout collections offer the ability to connect genotype to non-fitness related phenotypes through economical yet comprehensive screening and to test and inform models of microbial behavior. High-quality knockout collections made by transposon mutagenesis require the detailed annotation of extremely large transposon insertion mutant collections needed to achieve high coverage of complex microbial genomes (**Fig. 2**). However, the sophisticated hardware, time and cost needed to do this is out of reach for many investigators. Knockout Sudoku uses sophisticated reconstruction algorithms that allow for much simpler, faster and cheaper pooling, and leaves money in the budget for validation of the collection through orthogonal sequence analysis methods. The quality-controlled *S. oneidensis* Sudoku collection was assembled with a dedicated team of 2, for a budget of approximately #12,000, including validation and robot rental (**Supplementary Table 6**).

The knockout collection construction method presented here is generally applicable to any microorganism into which a transposon can be introduced. The ease of use and low cost of Knockout Sudoku opens the possibility of rapidly and cheaply constructing whole genome knockout collections for many esoteric organisms and even strains engineered for specific studies. These collections can facilitate comprehensive screening for the genes that underly many fascinating capabilities not seen in model organisms.

## Acknowledgements

We thank N. Ando, K. Davis, A. Palmer, S. Meisburger, B. Chang, E. Adler, K. Malzbender and C. Kyauk for experimental assistance; W. Metcalf for providing *E. coli* strain WM3064; J. Gralnick for providing *Shewanella oneidensis* MR-1; L. Kovacs, J. Miller, L.R. Parsons, S. Silverman, W. Wang and J. Wiggins for assistance with next generation sequencing and media preparation; N. Ando, P.A. Silver and J. Way for critical reading of this manuscript. This work was supported by a Career Award at the Scientific Interface from the Burroughs Wellcome Fund and Princeton University startup funds (B.B.).

## Author Contributions

B.B. and M.B. conceptualized and developed the Knockout Sudoku method and *S. oneidensis* knockout collection. B.B., M.B., L.S., I.A.A and O.A. designed and performed experiments. B.B. developed and implemented the analysis software. B.B. and M.B. wrote this article. All authors reviewed and revised the manuscript.

## Online Methods

### 1. Simulation of Transposon Insertion Mutant Collection Generation

The appropriate size of the progenitor collection was estimated by two methods: an analytical model based upon Poisson statistics that is a function solely of non-essential gene count, N, and a numerical Monte Carlo algorithm (**Supplementary Fig. 1, algorithm 1**) that combines the published genome sequence of *S. oneidensis* with previously published gene essentiality information.

The Monte Carlo algorithm generates a list of AT and TA dinucleotide^1^ positions that are suitable for transposon insertion in the *S. oneidensis* genome (GenBank accession numbers: chromosome: NC_004347.2; megaplasmid: NC_004349.1). The algorithm builds a lookup table indexed by possible insertion location, records the locus name in which this position falls and marks the location as either *“Dispensable*”, *“SurpriseDispensable”, “GuessDispensable”, “DispensableUnclearInEcoli”, “ExpectedEssential”, “NewEssential”* or *“Unknown”* using data from Deutschbauer *et al*.^*2*^. Possible insertion locations that did not fall into any locus were assigned to a non-coding locus and marked as *“Dispensable*”. We considered two scenarios: a low non-essential count, *N*, case where the disruptable genes were marked as *“Dispensable”; “SurpriseDispensable”; “GuessDispensable”* and *“DispensableUnclearInEcoli*”, and a high non-essential count case where all genes marked *“Unknown”* were also considered non-essential. The low non-essential count case considered 3,256 genes while the high case considered 4,184 genes.

Picking events were simulated by a random choice (with replacement) of a non-essential transposon insertion location. If the hit count for that locus was previously zero, the unique gene hit count, *n*_insertions_, was incremented by 1. In total, we computed 1,000 trials of 46,708 picks for both the high and low nonessential count scenarios and used these to compute the average value and standard deviation of *n*_insertions_ at each pick, *k* (**Fig. 2, solid lines**).

The size of a transposon insertion collection necessary to achieve adequate coverage of a microbial genome can also be calculated through an analytical model by assuming a Poisson distribution of transposon insertions. Assuming a genome with *N* non-essential genes, and a collection of *k* transposon insertion mutants, a given gene should see *x* insertions with a probability,

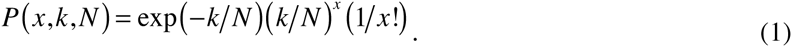

The probability that a single gene will see no transposon insertions,

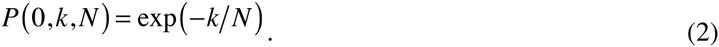

By linearity of expectation, the total number of genes that will see no insertions,

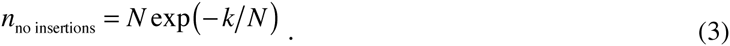

Or, the total number of genes for which insertions will be found,

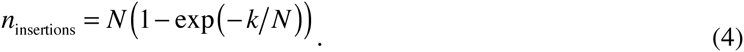

The analytically derived value of *n*_insertions_ is plotted for the high and low non-essential gene count scenarios in **Fig. 2** as dashed lines.

The simulated rarefaction curves were compared against an experimentally derived curve using a random sampling (without replacement) from the 46,708 non-unique transposon insertion instances in the progenitor collection catalog. This process was repeated 1,000 times to generate a standard deviation around the mean of the rarefaction curve (**Fig. 2, solid black lines**).

### 2. Preparation of Progenitor Collection

The progenitor transposon insertion collection was prepared by delivery of a modified pMiniHimar plasmid to *S. oneidensis* by conjugation with *E. coli* WM3064^3^. The transposon insertion library was plated onto LB agar Petri dishes with 30 μg/mL Kanamycin at a density of ≈ 200 colonies per plate. In total, 39,198 colonies were picked into the wells of 417 96-well polypropylene plates with 2 sterile control wells per plate with a CP7200 colony picking robot (Norgren Systems, Ronceverte WV, USA). Each well contained 100 μL per well of LB with 30 μg/mL Kanamycin. Plates were sealed with a sterile porous membrane (Aeraseal, Catalog Number BS-25, Excel Scientific) and incubated at 30 °C with shaking at 900 rpm until saturation was reached (typically ≈ 21 hours later).

### 3. Combinatorial Pooling and Cryopreservation of Progenitor Collection

The progenitor collection was pooled shortly after completion of growth using a 96-channel pipettor (Liquidator-96, Rainin) and speciality SBS format row (1 used) and column plates (1 used) (Catalog Numbers RRI-3028-ST and RRI-3029-ST, Phenix Research Products, Candler NC, USA) for row and column pool preparation and single well trays (OmniTray, Catalog Number 242811, Nalgene Nunc) for plate row (20 used) and plate column pool (21 used) preparation.

For each plate in the progenitor collection, 4 × 10 μL aliquots were withdrawn from each well with the 96-channel pipettor and dispatched to the row plate for row pooling; column plate for column pooling; a single-well plate designated as the plate-row pool receptacle for the plate; and another single-well plate designated as the plate-column pool receptacle for the plate. All 4 operations were completed in approximately 2 minutes. In total, the entire progenitor collection was pooled with 4 × 417 = 1,668 operations. Following pooling the plate was covered with a sterile lid and set aside until all plates were pooled.

Upon completion of pooling, all plates were cryopreserved by the addition of 50 μL (approximately equal to the remaining volume of culture after withdrawal of pooling aliquots and evaporation) of sterile 20% glycerol. Glycerol was mixed by shaking at 900 rpm and plates were immediately frozen at ‐80 °C. Pooling and cryopreservation was completed in a single day (≈ 18 hours) with a team of 5 people.

### 4. Pool Amplicon Library Generation and Sequencing

Amplicon sequencing libraries were generated from the mutant pools through a 2-step hemi-nested PCR reaction^4,5^. The reaction amplified a portion of the chromosome adjacent to the transposon present in each collection member and added Illumina TruSeq flow-cell-binding and read-primer-binding sequences to the 3’ and 5’ ends of the amplicon while replacing the standard Illumina index sequence with a custom barcode sequence for each pool. Genomic DNA was extracted from the coordinate pools with a genomic DNA extraction kit (Quick gDNA-MiniPrep, D3006, Zymo Research).

The first PCR step used a total reaction volume of 20 μL per well with 2 units of OneTaq DNA Polymerase (M0480, New England Biolabs, Ipswich MA, USA), 200 nM of each primer (Integrated DNA Technologies, Coralville IA, USA), and 200 μM of dNTPs (N0447, New England Biolabs). The first reaction was templated with 2 μL per well of purified genomic DNA at a concentration of ≈ 100 ng/μL, and used the three primers HimarSeq1.2, CEKG2C-IllR2, and CEKG2D-IllR2 (**Supplementary Table 4**) in the following reaction cycle: first 5 minutes at 95 °C; 6 cycles of 30 seconds at 95 °C, 30 seconds at 42°C (lowering by 1 °C per cycle), 3 minutes at 68 °C; then 24 cycles of 30 seconds at 95 °C, 30 seconds at 45 °C, 3 minutes at 68 °C; and finally 5 minutes at 68 °C followed by storage at 4 °C. Note that we do not attempt to normalize the amount of template loaded into the reaction for the amount of each species estimated to be in each pool.

Each second step PCR reaction used a total volume of 50 μL with 5 units of OneTaq polymerase and was templated with 0.5 μL of the completed corresponding first PCR reaction. The second step used 4 forward primers at 50 nM each (HimarSeq2.4-4N, HimarSeq2.4-5N, HimarSeq2.4-6N and HimarSeq2.4-7N).

Each of these primers contains between 4 and 7 random bases in order to avoid overloading a single color channel in the Illumina sequencer imaging system. Each amplicon pool used a different barcoding primer at 200 nM (**Supplementary Table 4**). This step used the reaction cycle: 30 cycles of 30 seconds at 95°C, 30 seconds at 56°C, 3 minutes at 68°C; 5 minutes at 68°C; and finally storage at 4 °C.

Products of this final reaction were pooled and were purified by agarose gel electrophoresis followed by gel extraction (Gel DNA Recovery Kit, Catalog Number D4001, Zymo Research) of the section of the library with molecular weights between 500 and 1000 bp. The samples were then sequenced on both lanes of an Illumina HiSeq 2500 in Rapid Run mode with 67 bp single end reads.

### 5. Analysis of Sequencing Dataset

We used a suite of custom algorithms developed in Python with the Scipy^6,7^ and Numpy^8^ libraries to condense the large volume of sequencing data into a collection address catalog (**Fig. 3**; **Supplementary Fig. 1**). Each read in the sequencing dataset was examined by a regular expression that matched the main sequence (Illumina read 1) with that of the transposon and the index sequence (Illumina read 2) to a pool barcode with up to 2 mismatches allowed in each case (**Supplementary Fig. 1, algorithm 2**). The genome sequence portions of those reads with a valid barcode and transposon sequence were aligned against the *S. oneidensis* MR-1 chromosome and megaplasmid sequences (GenBank ID accession numbers NC_004347.2 and NC_004349.1) using BOWTIE2^9^ in end-to-end alignment mode to yield a genomic transposon insertion coordinate. This data was condensed into a pool presence table that enumerates the number of reads corresponding to each pool for each transposon insertion coordinate, giving a set of coordinates that map back to the progenitor collection (**Fig. 3B**).

Each pool presence table line is thresholded to remove collection coordinates with low read counts. The read count threshold is scanned from 1 to 30 to determine a solution to the pool presence table in which the number of lines that map to single addresses is maximized while maintaining a large number of lines that map in any way and stabilizing the number that do not map. We found that a read count threshold of 5 produced the most satisfactory compromise between these requirements (**Supplementary Fig. 1, algorithm 3; Supplementary Fig. 3**).

Possible progenitor collection addresses were calculated for each line in the pool presence table by generating all possible combinations of coordinates with a read count above a predefined threshold. Lines with only one entry per coordinate axis mapped unambiguously to a single line, while lines with a multiple coordinates in only one axis mapped unambiguously to multiple lines.

A significant minority of lines map ambiguously to multiple addresses. An example of this is shown in line 3 of **Fig. 3B**. Intuitively, one would expect that if a given mutant in given well produced a large number of reads in one coordinate axis, it would also do so in the other 3. Thus, we expect that the 2 correct addresses for this line are composed of just high and just low sets of read count coordinates.

Pool presence table lines (transposon insertion coordinates) that map to single collection addresses are used to calculate the six ratios of reads between the pool axes (row/column; row/plate-column; row/plate row; column/plate-column; column/plate-row; plate row/plate-column). The natural logarithms of these ratios are used to generate a set of 6 histograms and are fit with a Voigt function (**Fig. 3C-H; Supplementary Fig. 1, algorithm 4**). The read number ratio fits were integrated and normalized to an area of 1.0 to generate a set of 6 probability distribution functions.

The read count ratios are calculated for each possible progenitor collection address that can be generated from the address coordinates in an ambiguously mapping pool presence table line. The probability of this read count ratio is assessed by using the probability distribution function for that ratio, and the total probability of the proposed address is calculated by multiplying these 6 probabilities together (**Supplementary Fig. 1, algorithm 5**). The most probable addresses, up to a maximum defined by the highest number of coordinates in a single axis for that line, are mapped to the collection catalog.

A small number of lines had coordinates in only 3 axes that were above threshold, but did have a coordinates in the 4^th^ axis. In these cases, the highest count coordinate was taken from the 4^th^ axis to make a complete set of coordinates and allow mapping.

A small number of transposon insertion locations produce amplicons that align to multiple consecutive coordinates in the *S. oneidensis* genome. These apparently consecutive mutants were grouped following mapping to locations in the progenitor collection. If the run of consecutive coordinates contains an AT or TA dinucleotide, the A of this pair is taken as the consensus coordinate. Otherwise, the sum of all reads used to map the mutants are used to calculate a median consensus coordinate.

### 6. Sanger Sequencing Verification of Location Inference Predictions

Mutant location inference predictions were verified by a 2-step hemi-nested PCR reaction^4,5^ similar to that used for pool amplicon library generation (**Online Methods 4**). The reaction amplified a portion of the chromosome adjacent to the transposon in the mutant.

The first PCR step used a total reaction volume of 20 μL per well with 2 units of OneTaq DNA Polymerase (M0480, New England Biolabs), 200 nM of each primer (Integrated DNA Technologies), and 200 μM of dNTPs (N0447, New England Biolabs). The first reaction was templated with 1 μL per well of saturated bacterial culture, and used the three primers HimarSeq2, CEKG2C and CEKG2D (**Supplementary Table 4**) in the following reaction cycle: first 5 minutes at 95 °C; 6 cycles of 30 seconds at 95 °C, 30 seconds at 42°C (lowering by 1 °C per cycle), 3 minutes at 68 °C; then 24 cycles of 30 seconds at 95 °C, 30 seconds at 45 °C, 3 minutes at 68 °C; and finally 5 minutes at 68 °C followed by storage at 4 °C.

Each second step PCR reaction used a total volume of 20 μL with 2 units of OneTaq polymerase, 200 nM of the primers HimarFRTSeq2 and CEKG4 (**Supplementary Table 4**) and was templated with 0.5 μL of the completed corresponding first PCR reaction. The following reaction cycle was used for the second step: 30 cycles of 30 seconds at 95°C, 30 seconds at 56°C, 3 minutes at 68°C; and finally 5 minutes at 68°C followed by storage at 4 °C.

The second step PCR reaction was purified by a standard PCR clean-up procedure (Genewiz, South Plainsfield NJ, USA) and sequenced by Sanger sequencing (Genewiz, South Plainsfield NJ, USA).

The measured transposon insertion location was found by alignment of the Sanger read against the *S. oneidensis* genome sequence with Blast^10^, and compared against the predicted transposon insertion location with a custom program developed with Python and the BioPython library^11^. A complete set of results is shown in **Supplementary Table 2**.

### 7. Selection of Mutants for Quality-Controlled Collection

The progenitor collection catalog (**Supplementary Table 1**) was used to assemble a quality-controlled non-redundant collection of mutants containing representatives for 3,282 genes disrupted in the progenitor collection.

We developed a set of algorithms to choose a representative mutant for each gene disrupted in the progenitor collection. Mutants were chosen for inclusion in the condensed collection by a scoring function that estimated the likelihood that a randomly picked colony struck out from a well containing a representative disruption of a gene would disrupt the function of that gene (**Supplementary Supplementary Fig. 4, algorithm 6**). The function first assessed the probability that the colony would be the desired mutant given the level of co-occupancy of the well. Then, given that the right colony was picked, the function estimated the probability that the disruption would knock out the function of the gene as the fractional distance of the transposon from its translation start.

We re-arrayed 2,699 mutants from singly-occupied wells and colony purified representatives for another 999 loci from multiply-occupied wells. For the mutants from multiply-occupied wells, we streaked a sample from each well and picked between 2 and 10 colonies, which we estimated would give an 85% chance of finding the desired mutant, into the wells of a series of 96-well plates. In total, this collection comprised of 5,653 samples, arrayed on 84 plates. The collection was arranged into a 10 × 9 plate grid, re-pooled and resequenced to assist in sorting well co-occupants from desired mutants (**Online Methods 4**).

### 8. Orthogonal Verification of Quality-Controlled Collection by Massively Parallel Sequencing

The sequencing data from the quality controlled-collection (**Online Methods 7**) was analyzed through an orthogonal method to test the sequence contents of each well against the predicted contents, rather than locate each sequence. The verification algorithm generates a list of all transposon coordinates appearing in each coordinate pool. The sequence content of all wells in the quality-controlled collection can be determined by calculating the intersection of the 4 transposon coordinate sets that correspond to the 4 pool coordinates of the well. If the intersection of the 4 pools contained one of the predicted genomic coordinates for that well, it was marked as correct. If the intersection contained the coordinate that we hoped to isolate by colony purification, it was also marked containing a desired mutant. The set-intersection algorithm was implemented in Python and the Scipy^6,7^ and Numpy^8^ scientific computing libraries.

### 9. *Shewanella* Basal Media

Our AQDS reduction assay experiments used a minimal media consisting of (per liter): ammonium chloride (NH_4_Cl) (0.46 grams); dibasic potassium phosphate (K_2_HPO_4_) (0.225 g); monobasic potassium phosphate (KH_2_PO_4_) (0.225 g); magnesium sulfate (MgSO_4_.7H_2_O) (0.117 g); ammonium sulfate ((NH_4_)_2_SO_4_) (0.225 g). The media was buffered with 100 mM HEPES its pH was adjusted to 7.2.

### 10. High-throughput Assay for Extracellular Electron Transfer Activity

To functionally validate the quality-controlled collection, we screened it for extracellular electron transfer activity with the AQDS screen developed by Newman and Kolter^12^. Briefly, the humic substance analog AQDS changes color from clear when oxidized to deep orange when reduced by 2 electrons (**Fig. 5A**).

A frozen stock of the quality-controlled *S. oneidensis* Sudoku collection was duplicated with a multi-blot replicator (Catalog Number VP 407, V&P Scientific, San Diego CA, USA) into 96-well plates containing 100 μL of LB media with 30 μg/mL Kanamycin per well. The plates were sealed with an air porous membrane (Aeraseal, Catalog Number BS-25, Excel Scientific) and grown to saturation overnight (typically 21 hours) at 30 °C with shaking at 900 rpm. The following day the plates were imaged with a photographic data acquisition system (the macroscope). An image analysis algorithm developed with Scikit Image^13^ and SimpleCV^14^ tested the plate images for cross-contamination and growth-failure events by comparison with the collection catalog. The image analysis algorithm updated the record for each well in the collection catalog with growth information to assist in rejection of false positives in the EET screen due to growth failure.

A 10 μL aliquot of culture from each well was transferred to a barcoded clear 96-well assay plate in which each well contained a mixture of 70 μL of buffered *Shewanella* Basal Media (**Online Methods 9**) and 20 μL of 25 mM AQDS, for a final concentration of 5 mM. The assay plates were immediately transferred to a vinyl anaerobic chamber (Coy Laboratory Products, Grass Lake MI, USA) with an atmospheric H_2_ concentration of ≈ 3%. Each plate in the collection was imaged multiple times over the course of approximately 48 hours with a macroscope device inside the anaerobic chamber. Hits were identified by visual inspection and an image analysis algorithm (**Online Methods 11**).

Hits were validated by colony purification and selection of 3 colonies into the wells of a 96-well plate. Re-test plates were grown to saturation overnight and tested again. Each re-test plate contained disruption mutants for genes flanking the candidate hits in the *S. oneidensis* genome that acted as quasi wild-type experimental controls and calibrants in image analysis. A representative sample was picked for each hit and its transposon identity was confirmed by Sanger sequencing.

### 11. Image Analysis

A custom image analysis program developed with SciKit Image^13^, SimpleCV^14^ and Matplotlib^15^ was used to sort the AQDS reduction assay images by barcode and identify well positions and assign well content information. Almost all information on the reduction state of the AQDS dye can be found in the blue color channel of the assay plate images. At the start of the assay, all color channels are saturated (resulting in a set of white wells). As the AQDS dye is reduced and becomes orange, the intensity of the red channel remains approximately constant, with a small reduction in green channel intensity and a large drop in the blue channel. Time courses of the reduction of the AQDS dye for each mutant (Supplementary **Fig. 5A** and **B**) were generated by calculating the mean blue channel intensity for all pixels at the center of each well in each image in the time series. Mutants that had grown successfully to saturation that displayed no significant change in blue channel intensity were reported as candidate hits.

The time series of colors for each gene shown as colored circles above each gene in **Figs. 5** and **6** and **Supplementary Fig. 6** were generated by an algorithm that interpolated the multi-replicate average of mean well-center color values for that mutant at 0, 10, 20 and 39.5 hours (the length of the shortest time series in all experiments) after the initiation of the reduction experiment.

Rates of reduction were calculated by a linear fit to the linear portion of the blue-channel reduction curve with Datagraph (Visual Data Tools). The reduction rate in units of moles per hour was computed using a conversion factor derived from the quasi wild-type controls. Results are shown in **Supplementary Table 5**.

